# When vision lags, motor prediction follows

**DOI:** 10.1101/2020.02.13.937235

**Authors:** Marieke Rohde, Gizem Altan, Marc O. Ernst

## Abstract

When we act, our brains register the visual feedback of the outcome of our actions only approximately 150 ms after sending the motor commands to the muscles. By anticipating the sensory consequences of our actions, temporal prediction mechanisms help us to accurately perform time-critical motor actions, such as catching a ball, and to experience the world as temporally coherent. It remains unclear whether these prediction mechanisms can adapt in a general way to changing sensory feedback delays, as they occur, for instance, when neural processing times increase during development. We here used Augmented Reality to train participants for one hour in a range of everyday life tasks with different levels of artificial visual feedback delay: 0 ms (No Delay), 100 ms (Small Delay) and 250 ms (Large Delay). We tested whether temporal aftereffects (increased motor-anticipation) generalize across different perceptual and motor tasks that participants did not practice during training and that involve different aspects of sensorimotor integration (motor planning, feedback control, sensorimotor time perception). Our results show a consistent temporal aftereffect of 33% across tasks and levels of delay. This suggests that the human sensorimotor system can adapt to the presence of visual feedback delays in a that affects sensorimotor prediction in general.

## Introduction

When first wearing newly prescribed glasses, we tend to feel dizzy due to unfamiliar geometrical distortions in our visual field. This dizziness typically vanishes within hours or days because our brain adapts by learning how changes in the visual field relate to our movement, and so restores the stability of action and perception^[1,2]^. Similar adaptation mechanisms should exist also to compensate for temporal distortions (delayed visual feedback): Even in natural settings, the brain must predict the consequences of our actions to compensate for processing delays between movement and vision that amount to an average of about 150 ms in human visuomotor control^[3]^, but that are variable depending on the stimulus or task.

Like with new glasses, the unexpected addition of small delays (e.g., 100 ms) to the visual feedback can interfere dramatically with motor coordination^[3-11]^ and perturbs the perceptual experience of action timing and causality^[3,5,11-13]^. To date, however, it is still unclear if the brain adapts to visual delays like it does to new glasses. That is, whether our sensorimotor prediction mechanisms can learn altered timing relations between action and feedback in a general manner that makes detrimental distortions vanish across perceptual and motor tasks when performed with a delay. Previous studies showed that when participants train to perform motor tasks with a visual delay, their motor performance improves^[3-11]^. However, it has been argued that this motor adaptation is merely of a task-specific nature^[7,9,10]^: The compensatory strategies acquired in these experiments help with the trained task and close variations, but there is little evidence that the adaptation effects generalize to other tasks^[5-11]^, as it would be expected if the visuomotor prediction mechanisms learned that an altered delay affects sensorimotor behavior in general.

We believe that in these earlier studies, the training procedures were too specific for the brain to adapt to the training delay in a general manner. We hypothesize that such general temporal adaptation occurs if participants are exposed to visual delays in the whole field of view that consistently occur when they perform many different movements for an extended period of time. Inspired by the classical sensorimotor adaptation studies using prism glasses^[1]^, we used Augmented Reality to expose participants to feedback delays in a broad range of real-world activities. We equipped participants with a set of stereo cameras in front of their eyes to record live pictures of their own actions from the ego-perspective. We directly relayed these recordings to them in a Head-Mounted-Display (VRMagic, Fig. 1a and Methods) with a delay of our choice. Participants thus saw the consequences of their actions with an added visual feedback delay.

**Figure 1.**
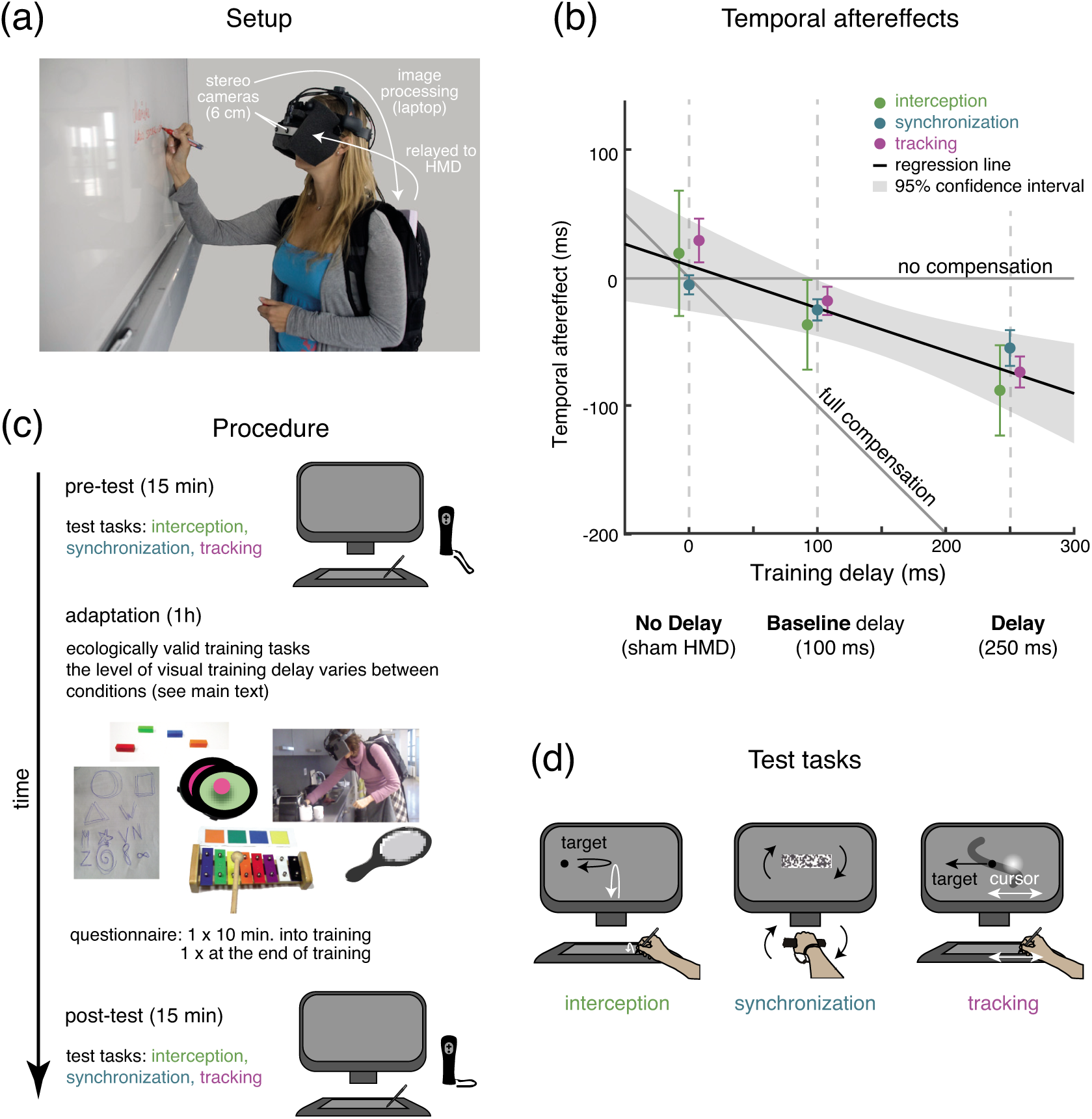
Procedure and Results (a) The Augmented Reality setup consists of a head-mounted display (HMD) with two cameras in front of the eyes (stereo-base of 6 cm; VRMagic Inc). Participants wore a backpack with a laptop to process the recorded images and relay them in real time to the HMD. (b) Main result: in all three test-tasks, temporal aftereffects of 33% of the training lag occurred (error bars show task means with standard error of the mean). (c) The procedure involved one hour of training in between pre- and post-tests in front of a computer (d) The test tasks: *Interception, Synchronization*, and *Tracking*.

Twenty participants were trained in Experiment 1 with a delay of 100 ms (*Small Delay* Condition). On a separate day, we retested the same 20 participants with a delay of 250 ms (*Large Delay* Condition). As a control, ten additional participants were trained without feedback delay: They wore a sham-setup consisting of goggles that restricted the field of view in a similar way as the headset (*No Delay* condition). During training, participants practiced a fixed sequence of everyday tasks for one hour with delay (Fig. 1c and Method Section). Before and after training, participants performed test tasks that were not part of the training procedure. We hypothesized that the training with feedback delays would cause a *temporal aftereffect*, such that participants would perform anticipatory control movements earlier after training with delays. To assess generalization, we tested for temporal aftereffects in three test tasks that emphasize different aspects of temporal processing for visuomotor control: The speeded, ballistic *Interception* task measured anticipation in motor planning. The *Synchronization* task measured perceived simultaneity of vision and hand movement. Finally, the continuous visually guided *Tracking* task measured temporal adaptation in feedback control.

## Results

Temporal aftereffects occurred in all three test tasks after the training with delays (Fig. 1b). The magnitude of the aftereffects is equal across tasks and scales linearly with the magnitude of the training delay (33% compensation of the training delay): A Linear Mixed Model analysis reveals a significant fixed effect of the slope r = −0.33 (confidence interval CI: [−0.53, −0.17]) but not of the intercept a = 10.2 ms (CI: [-20.5, 37.8] ms). A model comparison according to the Bayesian Information Criterion (BIC) favored this simple, two-parameter model over more complex models with additional fixed effects of task or with conditions coded as a nominal factor (see Table 1). This suggests that a single adaptation process underlies the adaptive changes observed with different levels of delay and in different tasks, i.e., that our neural visuomotor prediction mechanisms can adapt to altered sensorimotor timing (feedback delays) just as generally across tasks as it is known for spatial perturbations, such as prismatic distortions of the visual field.

**Table 1.** Model Comparison. All models have a random effect of the factor: Participant. Fixed Effects (FEs) modeled were: Task (T) with levels: *Interception, Synchronization* and *Tracking* (*Synchronization* and *Tracking* in Experiment 2 on *Intermanual Transfer*); Condition as continuous variable (CaC) with levels: 0 ms (*No Delay*), 100 ms (*Small Delay*), 250 ms (*Large Delay*); Condition as Factor (CaF) with levels: *No Delay, Small Delay* and *Large Delay* (*Large Delay* and *Intermanual Transfer* in Experiment 2 on *Intermanual Transfer*); x is an interaction, #par the number of parameters, BIC the Bayesian Information Criterion, AIC the Akaike Information Criterion and log-lik. the log-likelihood.

| ModelFEs | #par | BIC | AIC | log-lik. | deviance |
| --- | --- | --- | --- | --- | --- |
| **Experiment 1 on Delay Length** |  |  |  |  |  |
| CaC | 4 | **-242.48** | **-254.33** | 131.17 | -262.33 |
| CaF | 5 | -237.63 | -252.45 | 131.22 | -262.45 |
| CaC + T | 6 | -233.28 | -251.06 | 131.53 | -263.06 |
| CaF + T | 7 | -228.43 | -249.17 | 131.58 | -263.17 |
| CaC + T+ T x CaC | 8 | -224.71 | -248.41 | 132.2 | -264.41 |
| CaF + T+ T x CaF | 11 | -210.04 | -242.63 | 132.32 | -264.63 |
| **Experiment 2 on Intermanual Transfer** |  |  |  |  |  |
| CaF | 4 | **-148.08** | **-156.25** | 82.126 | -164.25 |
| CaF + T | 5 | -145.1 | -155.31 | 82.656 | -165.31 |
| CaF + T+ T x CaF | 6 | -141.52 | -153.78 | 82.888 | -165.78 |

To test the limits of motor generalization we conducted Experiment 2 asking whether there is *Intermanual Transfer* or whether adaptation is restricted to the effector used during training. Another ten participants were trained with the 250 ms visual feedback delay (*Large Delay*), undergoing the same training procedure as in Experiment 1, but with their right (dominant) hand immobilized. The untrained right hand was then used to perform the test-tasks. Even though the results are non-significant, they suggest a partial transfer of the temporal adaptation aftereffect from the non-dominant hand to the dominant hand (Fig. 2). Such a strongly reduced transfer effect across hands is in line with results from experiments on adaptation to spatial prismatic offsets of the visual field^[14]^.

**Figure 2.**
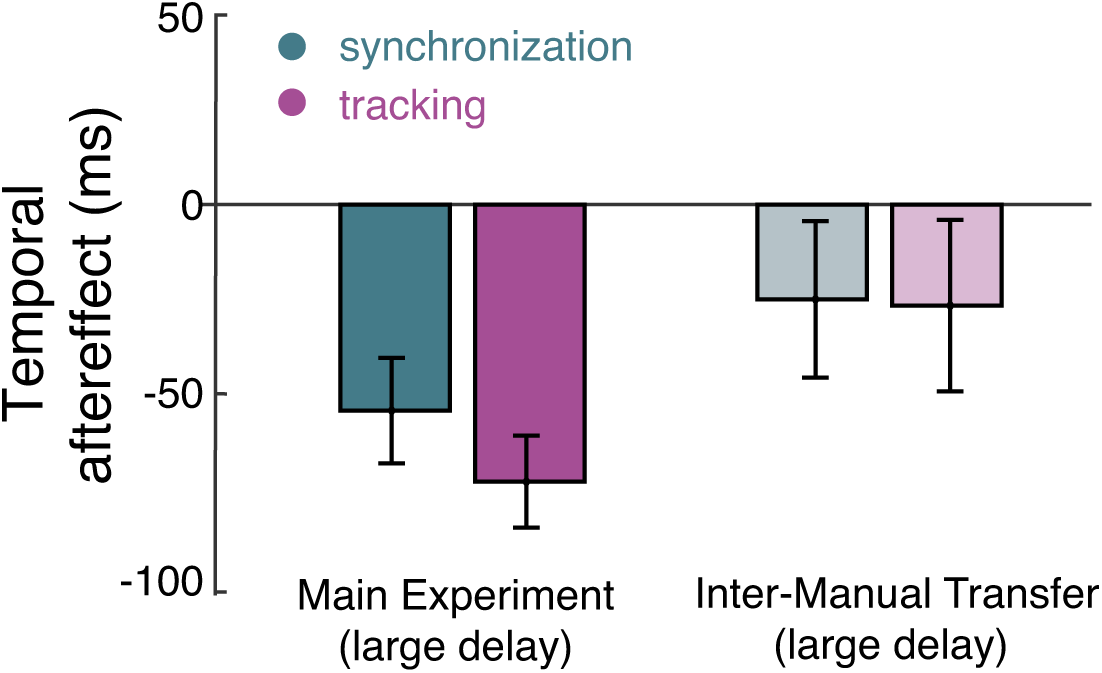
Intermanual Transfer. The *Synchronization* and *Tracking* test tasks were performed during the pre-test and post-test with the right, unexposed hand. The temporal aftereffect of −25.8 ms (10%, pale colors, CI: [-57.8, 5.1] ms) is not significantly different from either zero or the temporal aftereffect in the *Large Delay* condition of the Experiment 1 (dark colors; cf. Fig. 1). Error bars show task means with standard error of the mean.

## Discussion

In summary, the results show that prolonged exposure to delayed visual feedback leads to temporal adaptation of visuomotor prediction that generalizes across different domains of sensorimotor integration in untrained activities: motor planning, perceived visuomotor simultaneity, and feedback control. Moreover, upon debriefing, participants confirmed that they were not aware of the delay any more towards the end of training. That is, the training had restored their feeling of being in control of their body and actions. When first exposed to the 250 ms delay, some participants reported distortions in the perception of their own actions, such as “It is like the stick [that I move] is moving by its own”, “I know that I am initiating the movement, but it is not like my hand” or (in front of the mirror) “It does not feel like my hands belong to me”. Such reports are also known from the literature^[3,4,12,13]^. In one hour or less of training, all these subjective perturbations improved and many of them even disappeared. The transfer of delay adaptation across everyday-life tasks shows that our visuomotor delay compensation mechanisms are not hardwired to deal with fixed, naturally occurring delays. Delays in the sensorimotor loop can vary due to differences in neural processing time, physical causes such as ambient-light (or luminance), inertial lags, and more recently also due to transmission latencies in digital interfaces. The plasticity we report here allows humans to handle such variable delays in the sensorimotor loop and make their detrimental effects vanish across domains of sensorimotor processing for action and perception.

## Methods

### Participants

Fourty naïve participants took part in the study. Thirty participants took part in Experiment 1, out of which twenty participants (age range: 18 – 35 years, eleven female, two left-handed by self-report) were tested in both the *Large Delay* and *Small Delay* conditions (on separate days). Ten participants (age range: 21 – 31 years, five female, two left-handed by self-report) were tested in the *No Delay* control condition. Ten additional participants (age range, 21-26 years, eight female, all right-handed) took part in the Experiment 2 on *Intermanual Transfer* (one session with *Large Delay*, Fig. 2). The experiments were conducted in agreement with the ethics standards laid out in the Declaration of Helsinki and was approved by the ethics committee of the Department of Medicine of the University of Tübingen (Germany). All participants signed an informed consent form and received 6 €/h for their participation. The piloting participant depicted on Fig. 1 (a) has provided written informed consent for the publication of the image in an online open-access publication. The person visible in Fig. 1 (c) is one of the authors.

### Augmented Reality setup

For the *Small Delay* and *Large Delay* conditions, we used a digitally-mediated Augmented Reality headset (VRmagic GmbH, Mannheim, Germany) with a head-mounted display (HMD) and stereo cameras (VRmC-12+/C-COB USB, VRmagic GmbH, Mannheim, Germany). The cameras were mounted ca. 3cm in front of the eyes (angle of view: 40° horizontally, 25.4° vertically, frame rate: max. 70 Hz, exposure time: min. 30μs, resolution: 754 x 480) to record the visual scene. Custom-written C++ software that connected to the system’s API for input and output relayed the stereoscopic images from the cameras to the OLED displays in the HMD at 30 Hz. It was executed on a laptop that was carried by the participants in a backpack. The setup had a 98∓13 ms system delay (cf. 100 ms *Small Delay* condition). In the *Large Delay* condition, the images were delayed and replayed with an additional 150ms delay, i.e., the total delay was 250 ms. In the *No Delay* condition, participants were trained wearing a sham-setup consisting of analog goggles that restricted the field of view in a similar way as the HMD and the backpack with the unconnected experimental laptop.

### Test setup

Participants sat at a desk in front of a screen with their chin fixated at 63 cm viewing distance. All participants wore the headset with the baseline delay of 98∓13 ms during the test phase. The test stimuli were presented on a screen that participants saw through the headset. Participants’ hands were occluded from vision. Participants pressed the spacebar on a keyboard with their non-dominant hand to start test trials. They responded to the stimuli in the *Interception* and *Tracking* test-tasks using a stencil on a graphics tablet (WACOM Intuos 3 A3-wide; active area 48.8 cm × 30.5 cm and a grip pen; Wacom Europe GmbH, Krefeld, Germany). During the *Sychronization* trials, they responded with a Wii Remote controller (Nintendo Co., Ltd.). These responses were recorded using the open source DarwiinRemote software^[15]^. The room was kept dark during testing.

### Procedure

Before the first experimental session, participants came in for a short practice and screening procedure (<30 min). They were trained on the test tasks *(Interception*: 20 trials, *Synchronization*: 10 trials; *Tracking*: 20 trials). During the practice session, they were also given the questionnaire items to read for preparation and were instructed to ask for clarification if any of the questions were unclear to them.

Each experimental session lasted approximately 90 min and consisted of pre-test, training and post-test. Before the pre-test, participants performed two trials of each test tasks to refresh their memory. The pre-test and post-test lasted ca. 10-15 min each. The training lasted for one hour. The *Small Delay* and *Large Delay* conditions were recorded in the same population of participants but on different days. The starting condition was counter-balanced across participants. The *No Delay* control condition was recorded in a different participant population. Experiment 2 on *Intermanual Transfer* was recorded in a different participant population.

#### Test phase

During the test blocks (pre-test and post-test), participants sat at a desk wearing the HMD with the baseline lag of 98±13 ms (cf. Setup). Before the post-test started, participants were informed that the test would be performed under identical conditions as the pre-test and that any changes they might have experienced were now gone to avoid that participants would cognitively compensate for any delays they had become aware of. Pink noise was presented through earphones to cancel out auditory cues (e.g., stencil scratching on tablet). Participants performed the three test tasks (Fig. 1 (c)) in the following order: *Interceptio*n (seven trials), *Synchronization* (seven trials), and *Tracking* (five trials). All test tasks produced an estimate of signed *temporal error* (anticipatory movements were negative temporal error, lagging movements were positive temporal errors). All test tasks were implemented using the psychophysics toolbox for Matlab [16].

In *Interception* trials, a small black target disk (0.8° visual angle) entered from the left side of the participants’ field of view and decelerated towards the mid-line of the screen. Using a stencil on a graphics tablet, participants performed a speeded up-down movement (hand trajectory: 7.6 cm) to match the anticipated turning point of the disk in time. They were instructed to complete the movement within one second and were told that otherwise the trial would abort. The hand was occluded and there was no feedback about the movement beside a flashed red disk (0.8° visual angle; one frame) that was flashed halfway into the outward movement. Temporal error was defined as the difference in time between the turning point of the target and the turning point of the participant’s outward-inward hand movement.

In *Synchronization* trials, participants held a Nintendo Wiimote in the extended dominant hand. They saw a horizontal textured bar (height: 2.4° visual angle, width: 9.5° visual angle) on the screen that rotated by 45° clockwise and counterclockwise, following a sine-wave time course (the frequency of the rotation was drawn pseudo-randomly from [0.3 Hz, 0.4 Hz 0.5 Hz]; Fig. 1 (c)). Participants were instructed to rotate the Wiimote to match the movement of the bar such that the two movements felt simultaneous. There was no visual feedback and no feedback about whether participants performed well. The two time series (sign-inverted Wiimote acceleration, smoothed with a 10 Hz Butterworth filter, and target bar movement) were cross-correlated in a time window of ±500 ms, and the lag with the maximal cross-correlation was used as a measure of temporal error (cf. analysis of Exp. 3 in ^[11]^).

The *Tracking* task was a modified version of the tracking task used in^[11]^. Participants saw an irregular but continuous path (width 0.8° visual angle) scroll down from top to bottom. Their target was small black disk (0.8° visual angle) marking the middle of the path which moved irregularly to the left and right as the path scrolled down but remained at a fixed height (Fig. 1 (b); cf.^[11]^). Participants were instructed to track the movement of the target with a cursor by moving a stencil left and right on the graphics tablet. Their cursor was a white Gaussian blob with a standard deviation of 2.3° visual angle. The target movements extended up to 6° visual angle away from the midline, which corresponds to stencil movements of up to 3.2 cm away from the midline. The two time-series (stencil movement, smoothed with a 10 Hz Butterworth filter, and target movement) were cross-correlated in a time window of ±500 ms. The lag with the maximal cross-correlation was used as a measure of temporal error (cf.^[11]^).

#### Training phase

After completing the pre-test, participants sat with their eyes closed while the experimenter introduced the training delay and placed the laptop in the backpack. During the experiment on *Intermanual Transfer*, this preparation also involved the immobilization of the right hand by dressing participants in a jacket with a right glove sewn into the pocket. They inserted their right hand into this glove and were instructed not to remove their hand from the glove unless the experimenter told them to.

The training phase of the experiments consisted of seven main tasks and an optional eighth task.

1. Participants stood in front of a mirror (50 × 70 cm), where they could see the upper part of their body. They were instructed to execute movements, such as widely opening and then closing their arms horizontally and vertically, reaching, clapping (in the *Intermanual Transfer* experiment, they high-fived the experimenter instead) and observing their mirror image at the same time.
2. Participants navigated along lab corridors to a kitchen and back, guided by the experimenter. They were asked to explore their environment by touching or reaching the objects (e.g. reaching/touching door handles and columns). This was performed twice during the training (after tasks 1 and 6).
3. Participants completed everyday life tasks in the kitchen, following the experimenter’s instruction (taking a mug out of the cupboard, preparing a coffee for themselves or the experimenter, putting items in or out the dishwasher). This was performed both times that the participants navigated to the kitchen.
4. Participants played a commercially available buzz-wire game, where they had to move a hook on a handle along a curved wire, avoiding any contact between the hook and the wire. The task was to complete the path as fast as possible while avoiding contact with the wire. In case of a contact, the toy gave audiovisual feedback: a red light turned on for a brief time and a crash noise played. The duration of exposure to this task was from 5-10min and varied according to the performance and the speed of the participants.
5. Participants played sequences of tones on a toy xylophone with keys in different colors. The experimenter read out color sequences of three or four notes and the participants had to play these sequences rhythmically (i.e. keeping the time difference between each note the same) until the reproduction was deemed smooth by the experimenter. The entire task lasted 10-15min, depending on the performance and speed of the participants.
6. Participants were given a pen and a paper to trace the outline of geometrical shapes on paper (spirals, stars, flowers). Participants also traced the digits from 0 to 9 with a marker on a white board on the way back to experimental room. This task lasted 5min.
7. Participants had to slide, pick up and put down coins and colored wooden blocks on a table surface, following the beat of a metronome. These actions were performed repeatedly, starting with a rhythm of 30 beats per second. Once they improved performance, the rhythm was increased to up to 80 beats per second. The tasks lasted in total 10min.
8. If participants had completed the tasks extraordinarily fast, they were trained in an additional catching task. Participants were assisted to sit down on the laboratory room’s carpet. They visually pursued a tennis ball that rolled towards them and had to intercept the ball’s movement.

In both experiments and all conditions (*Large Delay, Small Delay, No Delay, Intermanual Transfer*) the tasks and instructions were the same and completed in the same order. The execution order of training tasks was: 1, 2, 3, 4, 5, 1, 2, 3, 6, 7, (8).

After completing the training, the experimenter removed the mobile setup and participants sat at the table to perform the post-test. Before starting the post-test, the experimenter informed the participants that the change that was introduced during the training was removed and their task is the same as before (i.e. pre-test). The participants were instructed to keep their eyes closed while being seated and waiting for the experimenter to prepare the setup.

### Questionnaire

We also applied a questionnaire at the beginning and at the end of the training phase during Experiment 1. The objective of this questionnaire was to record the changes in subjective experience that participants reported when the delay was introduced and when adaptation had taken place (see anecdotal results in main text). Even though the results captured some of the mentioned changes in subjective experience and did not contradict these reports, it was all in all not conclusive and did not go beyond what was already known from the literature. The results are therefore not reported here. The questionnaire was applied twice within each training phase; at the beginning (15 min after starting) and at the end (5min before ending) of training. Participants listened to the questionnaire items that were read out to them by a free online text-to-speech tool. They were asked to indicate the extent of their agreement or disagreement with twelve statements using a 7-point Likert scale.

### Data Analysis

Each trial of each test task yielded an estimate of temporal error (cf. description of test tasks above). The results were analyzed in Matlab 2015b, using functions of the Statistics Toolbox and the Signal Processing Toolbox.

*Temporal aftereffects* were defined as the change in temporal error between pre-test and post-test. The temporal aftereffect for each participant and condition was calculated as the across-trial median temporal aftereffect for the *Interception* and *Synchronization* tasks. For the *Tracking* task, the temporal aftereffect between the last pre-test trial and the first post-test trial was used for each participant and condition because of rapid re-adaptation under feedback when the delay was removed during the post-test (cf. ^[11]^). The population temporal aftereffects from Experiment 1 were modeled as a Linear Mixed Model (LMM) using R and the mixed modeling toolbox lme4^[17]^. In total seven data points of 150 were missing as a consequence of trial exclusion (see criteria below), i.e. for the respective participant, condition and task there was no valid data during either pre-test or the post-test. Different LMMs were compared according to the Bayesian Information Criterion (BIC). The simplest LMM had the lowest BIC, i.e., a model that regresses temporal error to training lag with just two fixed effects (one for the slope and one for the intercept) and one random effect (for participant). More complex models with additional fixed effects of task or with conditions coded as a nominal factor had a higher BIC (see Table 1). To compare the *Tracking* and *Synchronization* results from Experiment 2 on *Intermanual Transfer* with those of the *Large Delay* condition of Experiment 1, another LMM was fitted. Again, model comparison according to the BIC favored the simplest model, that is, a model with fixed effects for condition (as nominal factor) and a random effect for subjects (See Table 1).

### Data Exclusion

In all test tasks, trials were excluded according to specific criteria outlined in the following: *Interception.* In Experiment 1, trials were excluded if the absolute temporal error was larger than 500 ms (175 out of the 700 trials in Experiment 1). In Experiment 2 on *Intermanual Transfer*, the results from the *Interception* task had to be discarded entirely. The experimenter erroneously instructed participants to anticipate the movement of the target, not to intercept it. As a consequence, already in the pre-test, participants displayed an anticipatory temporal error of - 300±136 ms (mean and SEM), i.e., they did not perform the required task to intercept the target. Even before the training took place, this was significantly different from the pre-test temporal error of the *Large Delay* condition of the Experiment 1, which was −24±47 ms (mean and SEM; two sample t-test: t (58) = 2.3, p = 0.026).

#### Synchronization

Two pre-test *Synchronization* blocks are missing because the wiimote tracking device malfunctioned (This concerns one participant in the *Small Delay* condition and one participant in the *Large Delay* condition). Additionally, 166 out of the remaining 672 trials were excluded because the lag with the maximal cross-correlation was at the limit of the temporal window analyzed (i.e., −500 ms or +500 ms), which can indicate an artifact (the real maximum is outside the time window analyzed).

#### Tracking

24 out of the 500 *Tracking* trials were excluded because participants failed to perform the task (e.g., held their hand still). The exclusion criterion for these trials was defined in terms of tracking performance. Tracking error was defined as the squared distance between target and cursor integrated over a trial. The tracking error from all trials, participants and conditions was pooled and the outlier algorithm of the Matlab function boxplot.m was used to determine the exclusion threshold.

## Author contributions

MR and MOE developed the idea for the research. All authors developed the experimental setup and procedures. MR and GA conducted the measurements. MR and GA analyzed the data. All authors prepared the manuscript.

